# Cognitive and Motor Correlates of Grey and White Matter Pathology in Parkinson’s Disease

**DOI:** 10.1101/2020.05.04.077040

**Authors:** Mahsa Dadar, Myrlene Gee, Ashfaq Shuaib, Simon Duchesne, Richard Camicioli

**Affiliations:** Department of Radiology and Nuclear Medicine, Faculty of Medicine, Laval University; Department of Medicine, Division of Neurology, University of Alberta

**Keywords:** Parkinson’s disease, white matter hyperintensities, grey matter atrophy, deformation based morphometry, Scoring by Nonlocal Image Patch Estimator.

## Abstract

**Introduction:** Previous studies have found associations between grey matter atrophy and white matter hyperintensities (WMH) of vascular origin with cognitive and motor deficits in Parkinson’s disease (PD). Here we investigate these relationships in a sample of PD patients and age-matched healthy controls.

**Methods:** Data included 50 PD patients and 45 age-matched controls with T1-weighted and FLAIR scans at baseline, 18-months, and 36-months follow-up. Deformation-based morphometry was used to measure grey matter atrophy. SNIPE (Scoring by Nonlocal Image Patch Estimator) was used to measure Alzheimer’s disease-like textural patterns in the hippocampi. WMHs were segmented using T1-weighted and FLAIR images. The relationship between MRI features and clinical scores was assessed using mixed-effects models. The motor subscore of the Unified Parkinson’s Disease Rating Scale (UPDRSIII), number of steps in a walking trial, and Dementia Rating Scale (DRS) were used respectively as measures of motor function, gait, and cognition.

**Results:** Substantia nigra atrophy was significantly associated with motor deficits, with a greater impact in PDs (p<0.05). Hippocampal SNIPE scores were associated with cognitve decline in both PD and controls (p<0.01). WMH burden was significantly associated with cognitive decline and increased motor deficits in the PD group, and gait deficits in both PD and controls (p<0.03).

**Conclusion:** While substantia nigra atrophy and WMH burden were significantly associated with additional motor deficits, WMH burden and hippocampal atrophy were associated with cognitive deficits in PD patients. These results suggest an additive contribution of both grey and white matter damage to the motor and cognitive deficits in PD.

## INTRODUCTION

Parkinson’s disease (PD) is a progressive neurodegenerative disorder characterized by motor (bradykinesia, tremor, muscular rigidity, impaired postural reflexes, and gait dysfunction) and non-motor deficits (REM sleep behaviour disorder, autonomic dysfunction, neuropsychiatric symptoms, cognitive impairment and dementia)^1^. Two key pathological features define PD: degeneration of the nigrostriatal dopaminergic system and accumulation of misfolded α-synuclein aggregates spreading in a stereotypical caudal-to-rostral pattern^2^. Atrophy in the basal ganglia, particularly in the substantia nigra, subthalamic nucleus, nucleus accumbens, putamen, caudate nucleus, and internal and external globus pallidus, has been associated with motor and cognitive symptoms and longitudinal disease progression in PD^3,4^.

In addition to the typical PD pathology, co-morbid Alzheimer’s disease (AD) pathology has been observed in 20-30% of PD patients (with an even higher prevalence in patients with dementia)^2^ and atrophy in the hippocampi has also been related to slowing of gait and cognitive impairment in PD patients^5–8^.

White matter hyperintensities (WMHs) are areas of increased signal on T2-weighted or Fluid-attenuated inversion recovery (FLAIR) magnetic resonance imaging (MRI) sequences. WMHs of presumed vascular origin are common findings in the aging population and are attributed to an interplay of ischemic, inflammatory, and protein deposition processes. Hypertension and blood pressure fluctuations are associated with WMHs^9^. In non-PD elderly individuals, WMHs have been associated with global cognitive impairment, executive dysfunction, rigidity, and gait disorders^10–15^. Given that some of the symptoms associated with WMHs in non-PD individuals overlap with PD features, one might hypothesize that coexistence of WMHs in PD patients would lead to even greater cognitive and motor deficits. However, WMH associations with cognitive and motor symptoms are less well established and consistent in PD, although some recent studies report such relationships in both *de novo*^16^ and later stage PD patients^7,8,17–21^. Methodological differences in assessment of WMHs might explain some of these inconsistensies in the findings, where more recent studies use automated tools to measure WMH volumes^22,23^, while previous studies used semi-quantitative visual rating scales^24,25^, known to be less sensitive and more variable.

The aim of this study was to take advantage of our sample of PD and control participants for whom longitudinal MRI and clinical assessments were available, in order to investigate the impact of grey matter atrophy and textural changes, alongside WMH, on cognitive and motor symptoms in PD when compared to non-PD controls, as well as determine the strength of these relationships. We hypothesized that the greater the grey matter atrophy in substantia nigra regions and the hippocampus, together with a higher WMH burden, the greater the motor and cognitive deficits in PD patients when compared with age-matched normal controls.

## MATERIALS AND METHODS

### Subjects

Data included 50 subjects with PD (meeting UK brain bank criteria according to a movement disorders specialist) and 45 age-matched non-PD controls. Subjects were assessed at baseline, 18 and 36 months. Baseline age, sex, education and other demographic features were measured. As previously described^26^, general and neurological examinations were performed at each follow-up by a neurologist (RC), and included the Unified Parkinson’s Disease Rating Scale (UPDRS), a gait assessment (steps taken to traverse 9.1 meters), and the Dementia Rating Scale (DRS). Levodopa equivalent dose calculation and dementia assessment were performed as previously described^26^. Assessments of PD patients were performed in the ON state. Vascular risk factors were obtained at baseline. In brief, medical history and medications were reviewed for hypertension, diabetes, hyperlipidimia and ischemic disease at each visit. Postural vital signs (including supine and standing blood pressure) were obtained at each assessment. Blood was obtained at baseline and included measurement of fasting glucose and lipids as well as homocysteine, and Apolipoprotein E genotyping at the clinical laboratory at the University of Alberta Hospital^27,28^. The study was approved by the Health Research Ethics Board of the University of Alberta Pro00001182.

### MRI Acquisition

All participants received MRI scans on a Siemens Sonata 1.5T system. T1-weighted images were acquired using a 3D magnetization prepared rapid acquisition gradient echo sequence (MPRAGE) (TR=1800 ms, TE=3.2ms, TI=1100ms, 1 average, flip angle = 15°, field of view (FOV) = 256 mm, image matrix = 256 × 256, 128 slices, 1.5mm slices). Native spatial resolution was 1 × 1 × 1.5 mm^3^ which was zero-filled to 0.5 × 0.5 × 1.5 mm^3^. Axial FLAIR images were also acquired (TR = 8000 ms, TE = 99 ms, 2 averages, FOV = 220 mm, 25 slices, 5 mm slice thickness) oriented to the inferior margin of the corpus callosum.

### MRI Preprocessing

All T1-weighted and FLAIR images were preprocessed in three steps: noise reduction^29^, intensity non-uniformity correction^30^, and intensity normalization into range [0-100]. Using a 6-parameter rigid registration, the FLAIR and T1-weighted images were linearly coregistered. The T1-weighted images were first linearly^31^ and then nonlinearly^32^ registered to the MNI-ICBM152 average template^33^.

### Grey Matter DBM Measurements

Deformation based morphometry (DBM) maps were calculated by computing the Jacobian determinant of the deformation fields obtained from the nonlinear transformations to the average template. An atlas of cortical grey matter regions was used to calculate average grey matter atrophy in 80 cortical regions^33^. Similarly, an atlas of deep grey matter regions was used to calculate average subcortical grey matter atrophy in 16 subcortical regions^34^ (e.g. Figure 1.a. for substantia nigra).

**Figure 1.**
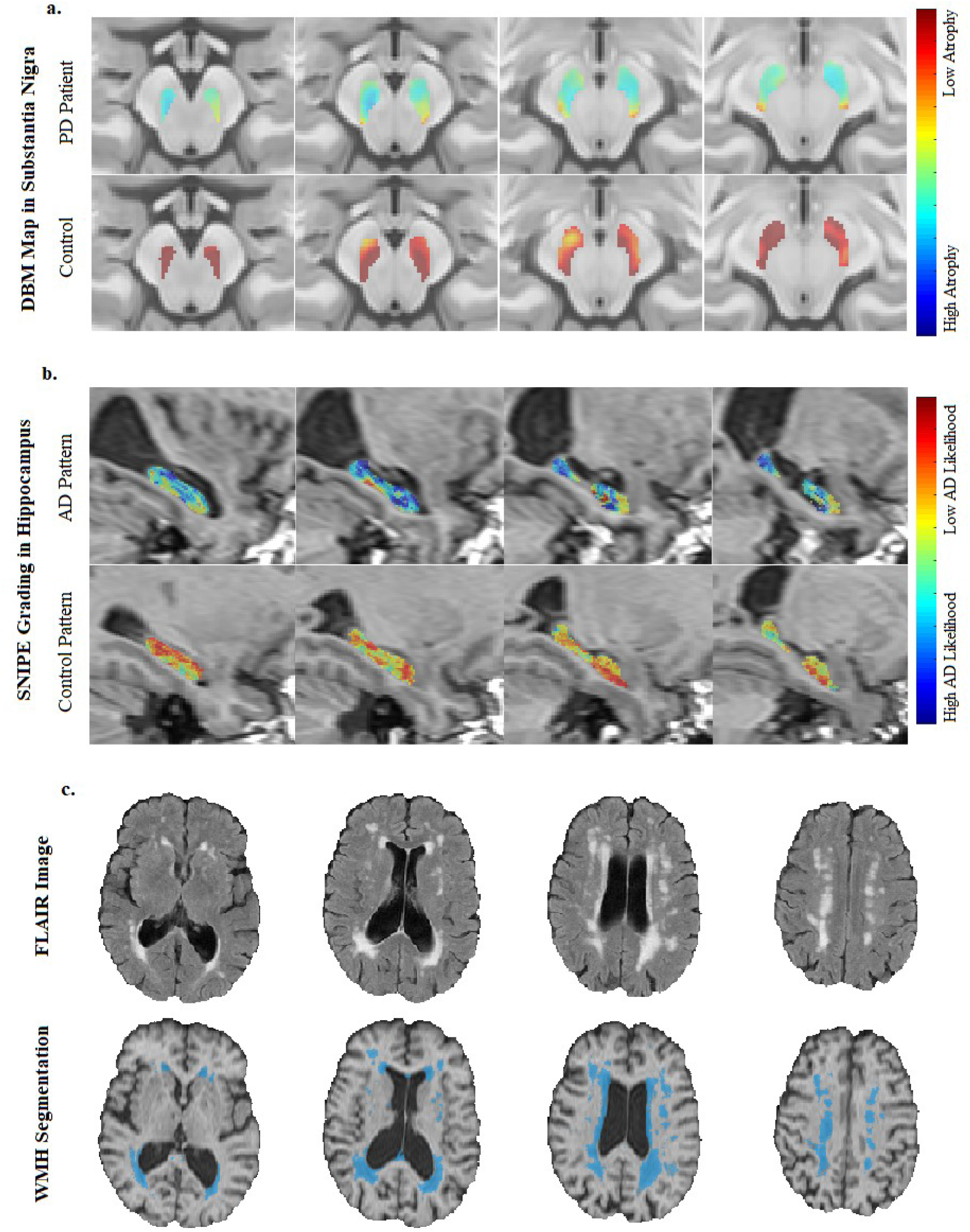
Extracted MRI Measurements: a. DBM maps in the substantia nigra mask for a PD and a control participant, b. hippocampal SNIPE grading for an AD-like and a control-like participant, and c. axial FLAIR scans and corresponding WMH segmentations. MRI= Magnetic Resonance Imaging. FLAIR= FLuid-Attenuated Inversion Recovery. WMH= White Matter Hyperintensity. DBM= Deformation Based Morphometry. PD= Parkinson’s Disease. SNIPE= Scoring by Nonlocal Image Patch Estimator. AD= Alzheimer’s Disease.

### SNIPE Grading in the hippocampi

An estimate of the level of pathology related to Alzheimer’s disease was obtained by applying Scoring by Nonlocal Image Patch Estimator (SNIPE) to the preprocessed T1-weighted images for all timepoints^35^. SNIPE assigns a similarity metric to each voxel in the hippocampus, indicating how much that voxel’s surrounding patch resembles similar patches in a library of patients with probable Alzheimer’s disease or cognitively healthy controls (Figure 1.b). The final SNIPE grading score is an average of all the voxels inside the hippocampal region, providing an estimate of the confidence that this image is similar to those of individuals with probable Alzheimer’s disease. Positive SNIPE grading scores indicate normal appearing hippocampi, whereas negate scores indicate presence of AD-like atrophy.

### WMH Measurements

The WMHs were segmented using a previously validated automatic multi-modality segmentation technique that combines a set of location and intensity features obtained from a library of manually segmented scans with a Random Forest classifier to detect WMHs^22,36^ (Figure 1.c). The library used in this study was generated based on data from the *Alzheimer’s Disease Neuroimaging* Initiative (ADNI) since the FLAIR images had similar acquisition protocols. WMH load was defined as the volume of the voxels identified as WMH in the standard space (in cm^3^) and are thus normalized for head size. WMH volumes were log-transformed to achieve normal distribution.

### Quality Assessment

The quality of all the registrations and segmentations were visually assessed and cases that did not pass this quality control were discarded (N=3, due to acquisition artifacts such as motion). All MRI processing, segmentation and quality control steps were blinded to clinical outcomes.

### Statistical Analyses

Paired t-tests or Chi-squared tests were used to assess differences between PD patients and controls in the baseline demographics and clinical variables. Mixed-effects models were used to investigate group differences in the MRI measurements (*eq.1*) as well as the associations between longitudinal MRI and clinical variables (*eq.2*). To assess the relationship between change in the MRI measurements during the time of the study and a diagnosis of dementia at the last visit (*m36*) in the PD group, age was split into age at baseline (*AgeBL*) and time from the baseline (*TimeFromBL*) visit (*eq. 3*).

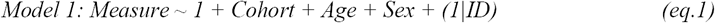

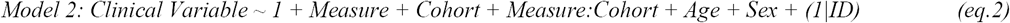

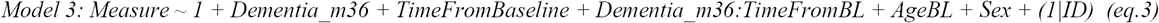

where *Cohort* denotes a categorical variable contrasting PD patients and controls, and *Measure:Cohort* denotes an interaction term between the measure of interest and *Cohort*, reflecting differences in the slope of change in the variable of interest between the two groups. *Dementia_m36* denotes a categorical variable contrasting a diagnosis of dementia at 36 months (i.e. last visit). *Clinical Variable* denotes the clinical variable of interest such as DRS or UPDRSIII scores. Subject ID was entered as a categorical random variable. To control for the impact of PD medications on motor symptoms, models assessing motor performance (i.e. UPDRSIII and gait measurements) also included Levodopa equivalent dose as a continuous random variable. All continuous variables were z-scored prior to the analyses. All atlas based results were corrected for multiple comparisons using False Discovery Threshold (FDR) technique with a significance threshold of 0.05.

## RESULTS

Table 1 summarizes the baseline demographics and clinical characteristics of the controls and PD patients used in this study. There were no significant differences in demographic variables. Compared to controls, PD patients had significantly lower standing systolic blood pressure (p=0.01) and greater drop in both systolic and diastolic blood pressure (p<0.002), lower total and LDL cholesterol levels (p<0.04), and significantly higher homocysteine levels (p<0.0001). As expected, PD patients also had significantly higher UPDRSIII scores (p<0.0001) and step count in the gait trial (p=0.05) than the controls. There were no significant differences in demographic variables between male and female participants. Compared to the female participants, the male participants had significantly lower HDL cholesterol levels in both control and PD groups (p<0.002). The control male participants also had significantly lower LDL and total cholesterol levels compared to the female control participants (p<0.01). There were no significant differences between male and female participants in other clinical characteristics in Table 1.

**Table 1.**
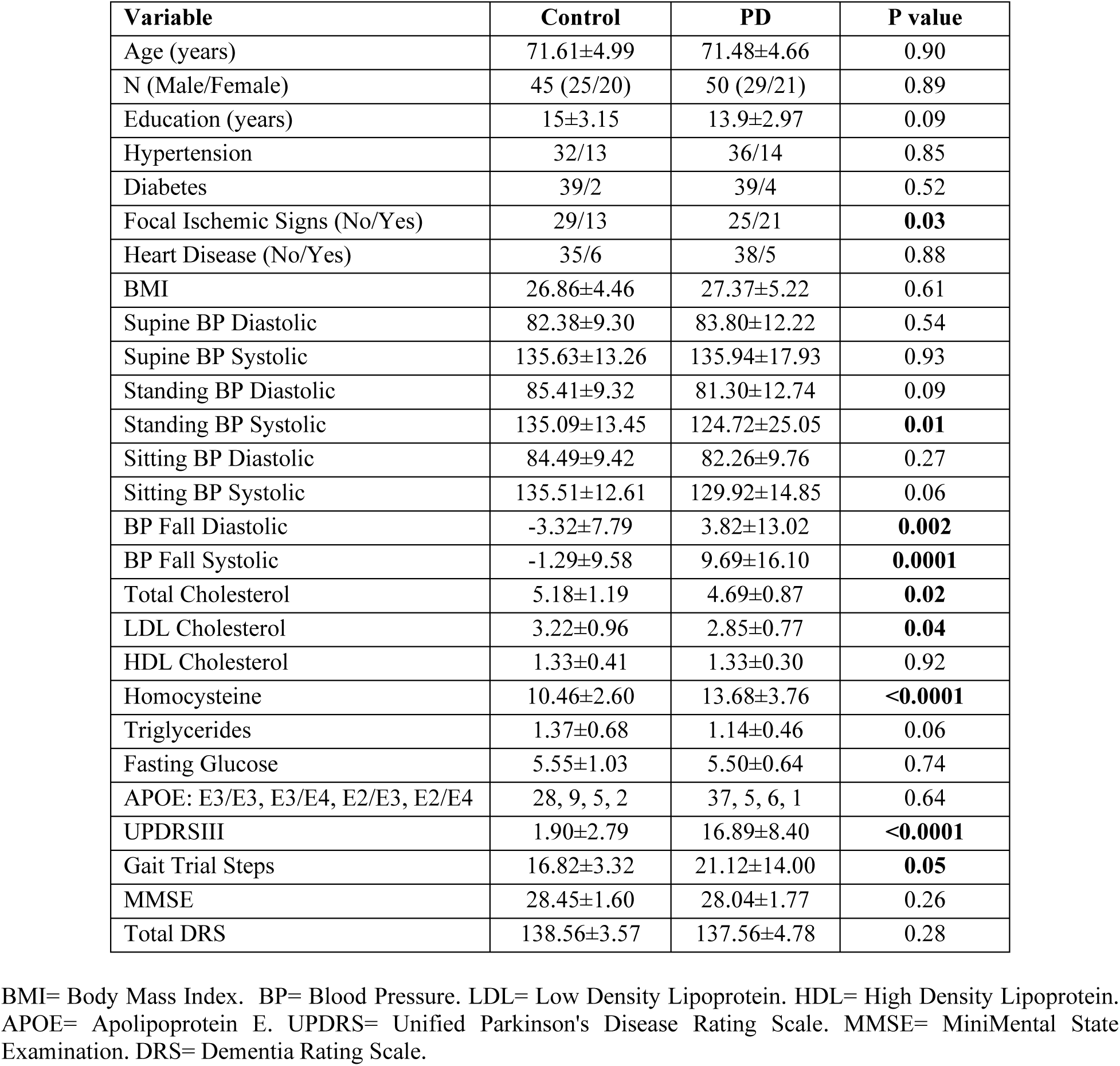
Descriptive statistics for the subjects enrolled in this study. Data are numbers (N) or mean ± standard deviation. P<0.05 is indicated in bold font.

| Variable | Control | PD | P value |
| --- | --- | --- | --- |
| Age (years) | 71.61±4.99 | 71.48±4.66 | 0.90 |
| N (Male/Female) | 45 (25/20) | 50 (29/21) | 0.89 |
| Education (years) | 15±3.15 | 13.9±2.97 | 0.09 |
| Hypertension | 32/13 | 36/14 | 0.85 |
| Diabetes | 39/2 | 39/4 | 0.52 |
| Focal Ischemic Signs (No/Yes) | 29/13 | 25/21 | **0.03** |
| Heart Disease (No/Yes) | 35/6 | 38/5 | 0.88 |
| BMI | 26.86±4.46 | 27.37±5.22 | 0.61 |
| Supine BP Diastolic | 82.38±9.30 | 83.80±12.22 | 0.54 |
| Supine BP Systolic | 135.63±13.26 | 135.94±17.93 | 0.93 |
| Standing BP Diastolic | 85.41±9.32 | 81.30±12.74 | 0.09 |
| Standing BP Systolic | 135.09±13.45 | 124.72±25.05 | **0.01** |
| Sitting BP Diastolic | 84.49±9.42 | 82.26±9.76 | 0.27 |
| Sitting BP Systolic | 135.51±12.61 | 129.92±14.85 | 0.06 |
| BP Fall Diastolic | -3.32±7.79 | 3.82±13.02 | **0.002** |
| BP Fall Systolic | -1.29±9.58 | 9.69±16.10 | **0.0001** |
| Total Cholesterol | 5.18±1.19 | 4.69±0.87 | **0.02** |
| LDL Cholesterol | 3.22±0.96 | 2.85±0.77 | **0.04** |
| HDL Cholesterol | 1.33±0.41 | 1.33±0.30 | 0.92 |
| Homocysteine | 10.46±2.60 | 13.68±3.76 | **<0.0001** |
| Triglycerides | 1.37±0.68 | 1.14±0.46 | 0.06 |
| Fasting Glucose | 5.55±1.03 | 5.50±0.64 | 0.74 |
| APOE: E3/E3, E3/E4, E2/E3, E2/E4 | 28, 9, 5, 2 | 37, 5, 6, 1 | 0.64 |
| UPDRSIII | 1.90±2.79 | 16.89±8.40 | **<0.0001** |
| Gait Trial Steps | 16.82±3.32 | 21.12±14.00 | **0.05** |
| MMSE | 28.45±1.60 | 28.04±1.77 | 0.26 |
| Total DRS | 138.56±3.57 | 137.56±4.78 | 0.28 |

BMI= Body Mass Index. BP= Blood Pressure. LDL= Low Density Lipoprotein. HDL= High Density Lipoprotein. APOE= Apolipoprotein E. UPDRS= Unified Parkinson's Disease Rating Scale. MMSE= MiniMental State Examination. DRS= Dementia Rating Scale.

### Vascular Risk Factors

Table 2 shows the associations between vascular risk factors and the MRI measures in controls and PD patients, controlling for age and sex. To decrease the number of analyses, left and right hippocampal SNIPE scores and substantia nigra DBM values were averaged. For each risk factor, use of relevant medication was included as a binary categorical covariate in the model (e.g. blood pressure medication for blood pressure measurements, diabetes medication for fasting glucose, cholesterol medication for cholesterol measurements, etc.) to ensure that the results are not driven by the medication usage.

**Table 2.**
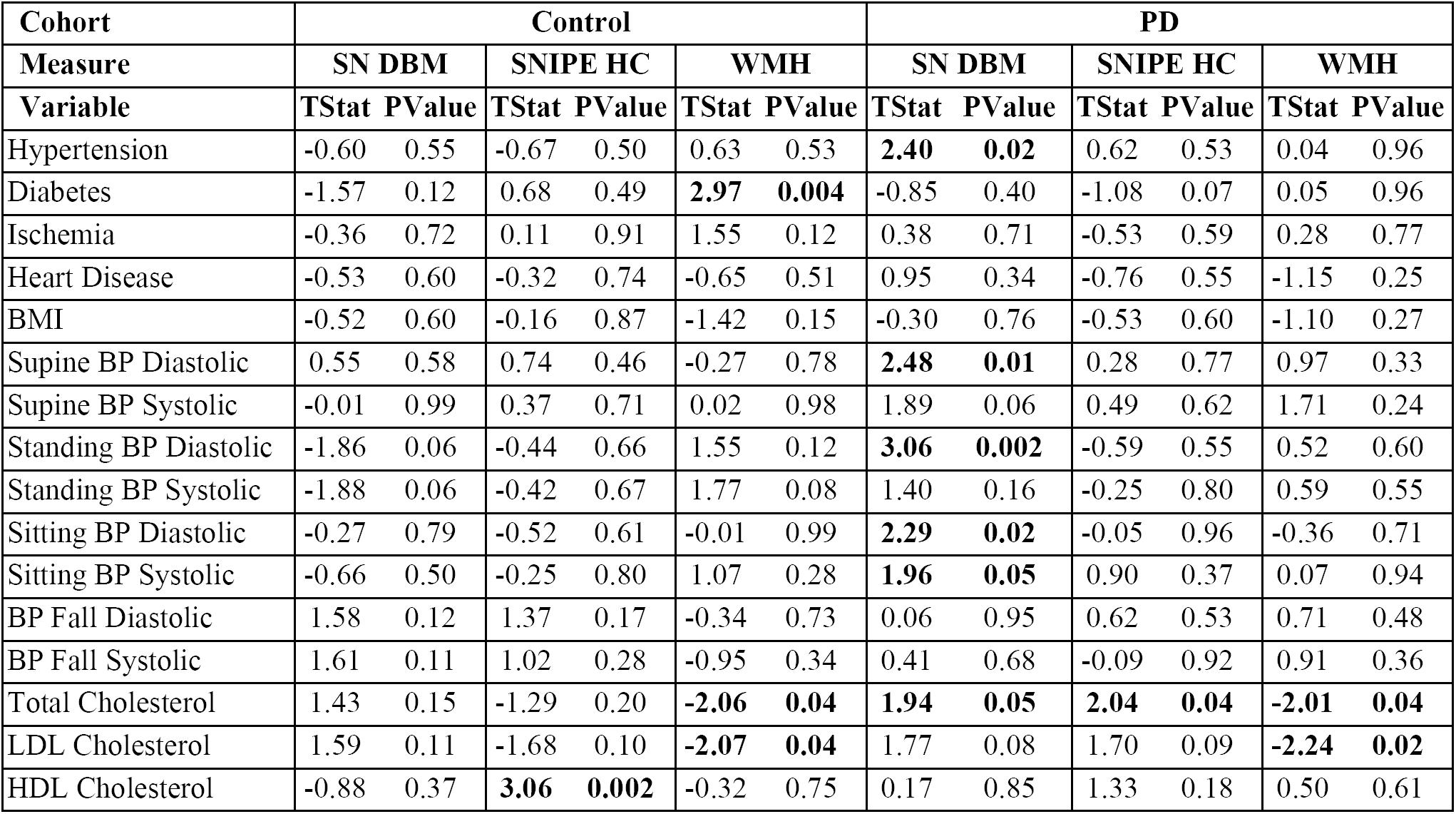
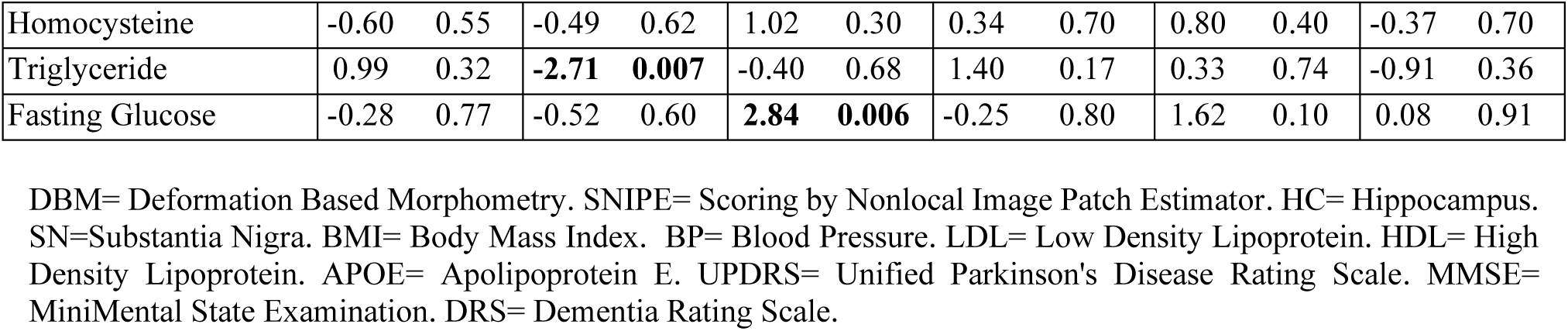
Association between substantia nigra DBM, SNIPE grading, and WMH load and vascular risk factors. Data are numbers (N) or mean ± standard deviation. P<0.05 is indicated in bold font.

| Cohort | Control |  |  |  |  |  | PD |  |  |  |  |  |
| --- | --- | --- | --- | --- | --- | --- | --- | --- | --- | --- | --- | --- |
| Measure | SN DBM |  | SNIPE HC |  | WMH |  | SN DBM |  | SNIPE HC |  | WMH |  |
| Variable | TStat | PValue | TStat | PValue | TStat | PValue | TStat | PValue | TStat | PValue | TStat | PValue |
| Hypertension | -0.60 | 0.55 | -0.67 | 0.50 | 0.63 | 0.53 | **2.40** | **0.02** | 0.62 | 0.53 | 0.04 | 0.96 |
| Diabetes | -1.57 | 0.12 | 0.68 | 0.49 | **2.97** | **0.004** | -0.85 | 0.40 | -1.08 | 0.07 | 0.05 | 0.96 |
| Ischemia | -0.36 | 0.72 | 0.11 | 0.91 | 1.55 | 0.12 | 0.38 | 0.71 | -0.53 | 0.59 | 0.28 | 0.77 |
| Heart Disease | -0.53 | 0.60 | -0.32 | 0.74 | -0.65 | 0.51 | 0.95 | 0.34 | -0.76 | 0.55 | -1.15 | 0.25 |
| BMI | -0.52 | 0.60 | -0.16 | 0.87 | -1.42 | 0.15 | -0.30 | 0.76 | -0.53 | 0.60 | -1.10 | 0.27 |
| Supine BP Diastolic | 0.55 | 0.58 | 0.74 | 0.46 | -0.27 | 0.78 | **2.48** | **0.01** | 0.28 | 0.77 | 0.97 | 0.33 |
| Supine BP Systolic | -0.01 | 0.99 | 0.37 | 0.71 | 0.02 | 0.98 | 1.89 | 0.06 | 0.49 | 0.62 | 1.71 | 0.24 |
| Standing BP Diastolic | -1.86 | 0.06 | -0.44 | 0.66 | 1.55 | 0.12 | **3.06** | **0.002** | -0.59 | 0.55 | 0.52 | 0.60 |
| Standing BP Systolic | -1.88 | 0.06 | -0.42 | 0.67 | 1.77 | 0.08 | 1.40 | 0.16 | -0.25 | 0.80 | 0.59 | 0.55 |
| Sitting BP Diastolic | -0.27 | 0.79 | -0.52 | 0.61 | -0.01 | 0.99 | **2.29** | **0.02** | -0.05 | 0.96 | -0.36 | 0.71 |
| Sitting BP Systolic | -0.66 | 0.50 | -0.25 | 0.80 | 1.07 | 0.28 | **1.96** | **0.05** | 0.90 | 0.37 | 0.07 | 0.94 |
| BP Fall Diastolic | 1.58 | 0.12 | 1.37 | 0.17 | -0.34 | 0.73 | 0.06 | 0.95 | 0.62 | 0.53 | 0.71 | 0.48 |
| BP Fall Systolic | 1.61 | 0.11 | 1.02 | 0.28 | -0.95 | 0.34 | 0.41 | 0.68 | -0.09 | 0.92 | 0.91 | 0.36 |
| Total Cholesterol | 1.43 | 0.15 | -1.29 | 0.20 | **-2.06** | **0.04** | **1.94** | **0.05** | **2.04** | **0.04** | **-2.01** | **0.04** |
| LDL Cholesterol | 1.59 | 0.11 | -1.68 | 0.10 | **-2.07** | **0.04** | 1.77 | 0.08 | 1.70 | 0.09 | **-2.24** | **0.02** |
| HDL Cholesterol | -0.88 | 0.37 | **3.06** | **0.002** | -0.32 | 0.75 | 0.17 | 0.85 | 1.33 | 0.18 | 0.50 | 0.61 |
| Homocysteine | -0.60 | 0.55 | -0.49 | 0.62 | 1.02 | 0.30 | 0.34 | 0.70 | 0.80 | 0.40 | -0.37 | 0.70 |
| Triglyceride | 0.99 | 0.32 | **-2.71** | **0.007** | -0.40 | 0.68 | 1.40 | 0.17 | 0.33 | 0.74 | -0.91 | 0.36 |
| Fasting Glucose | -0.28 | 0.77 | -0.52 | 0.60 | **2.84** | **0.006** | -0.25 | 0.80 | 1.62 | 0.10 | 0.08 | 0.91 |

DBM= Deformation Based Morphometry. SNIPE= Scoring by Nonlocal Image Patch Estimator. HC= Hippocampus. SN=Substantia Nigra. BMI= Body Mass Index. BP= Blood Pressure. LDL= Low Density Lipoprotein. HDL= High Density Lipoprotein. APOE= Apolipoprotein E. UPDRS= Unified Parkinson's Disease Rating Scale. MMSE= MiniMental State Examination. DRS= Dementia Rating Scale.

Substantia nigra DBM scores were not associated with any of the risk factors in the controls. In the PD group, hypertension, higher diastolic blood pressure (in all supine, standing, and sitting positions), higher systolic blood pression in a sitting position, and higher total cholesterol levels were associated with higher DBM scores in the substantia nigra (p<0.05). In the control group, higher HDL cholesterol levels and lower triglyceride levels were associated with higher hippocampal SNIPE scores, indicating lower levels of AD-related atrophy (p<0.007). In the PD group, higher total and LDL cholesterol levels were associated with higher hippocampal SNIPE scores (p<0.04). The control participants with a history of diabetes and higher fasting glucose levels had significantly greater WMH loads (p<0.006). In both controls and PD patients, higher levels of total cholesterol and LDL cholesterol were associated with lower WMH volumes (p<0.04). There were no significant differences between male and female participants in either group.

### Group Comparisons

After FDR correction, mean DBM values were significantly lower in PD patients compared to controls in left (t=-3.37, uncorrected p=0.0001, Figure 2.a) and right accumbens areas (t=-3.18, uncorrected p=0.001, Figure 2.b), left (t=-4.19, uncorrected p<0.0001, Figure 2.c) and right (t=-3.95, uncorrected p=0.0001, Figure 2.d) substantia nigra, and right subthalamic nucleus (t=-3.23, uncorrected p=0.001, Figure 2.e). Mean DBM values in left and right substantia nigra and right subthalamic nucleus also significantly decreased with age in both controls and PD patients (uncorrected p<0.0001). In both controls and PD patients, WMH burden significantly increased with age (t=7.71, p<0.0001, Figure 2.e). There were no significant group differences in WMH loads (t=0.49, p=0.62). There were no significant sex differences in the DBM and WMH measurements in either group.

**Figure 2.**
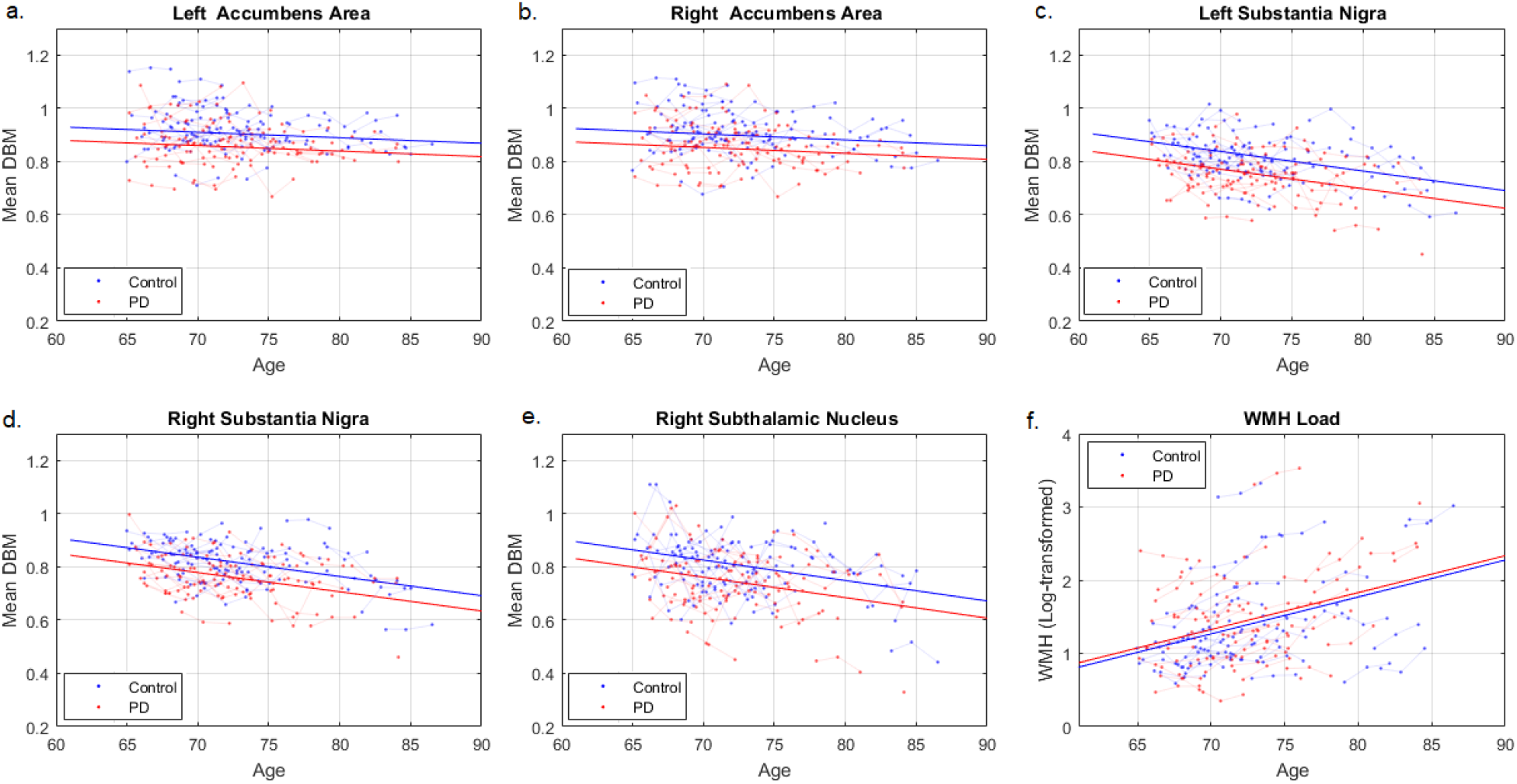
Group differences in average grey matter DBM values and WMH load. DBM= Deformation Based Morphometry. PD= Parkinson’s Disease. WMH= White Matter Hyperintensities.

### Presence of dementia at 36 months

Controlling for age at baseline and sex, WMH load significantly increased in PD patients that were diagnosed with dementia at 36 months (t=3.58, p=0.0005), compared to those that remained stable (Figure 3.a). There was also a marginally significant decrease in the right hippocampal SNIPE grading scores (t=-1.69, p=0.09, Figure 4.c). There were no significant sex differences in WMH and SNIPE measurements of the patients that are diagnosed with dementia at 36 months and those that are not.

**Figure 3.**
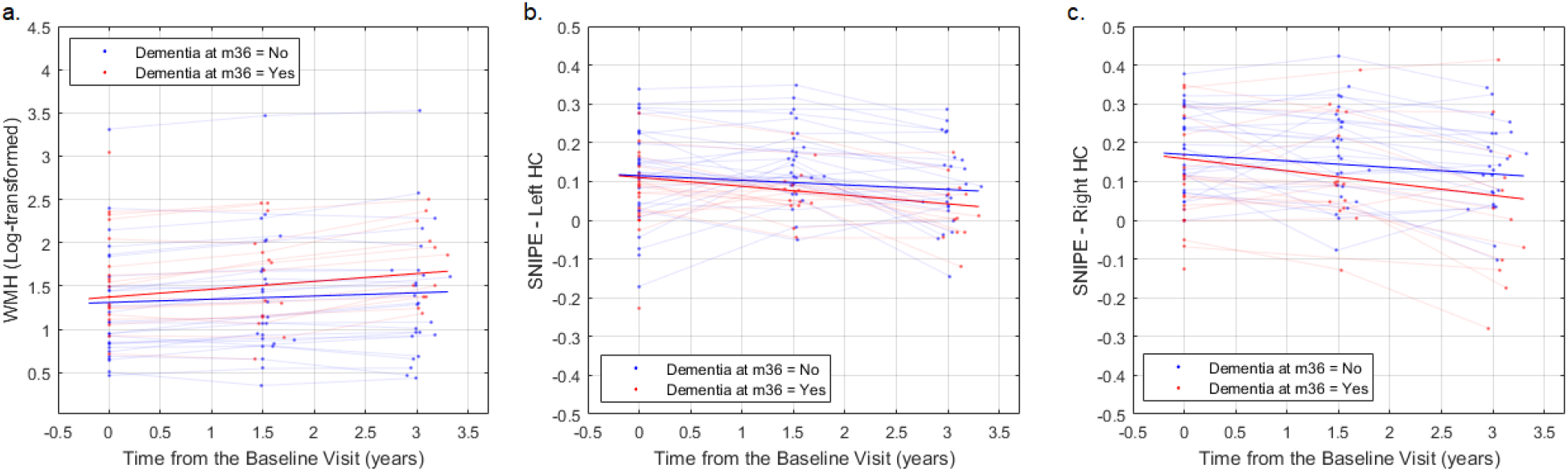
Comparing the change in the WMH load and the hippocampal SNIPE grading scores in PD patients that are diagnosed with dementia at 36 months and those that remain stable. SNIPE= Scoring by Nonlocal Image Patch Estimator.

**Figure 4.**
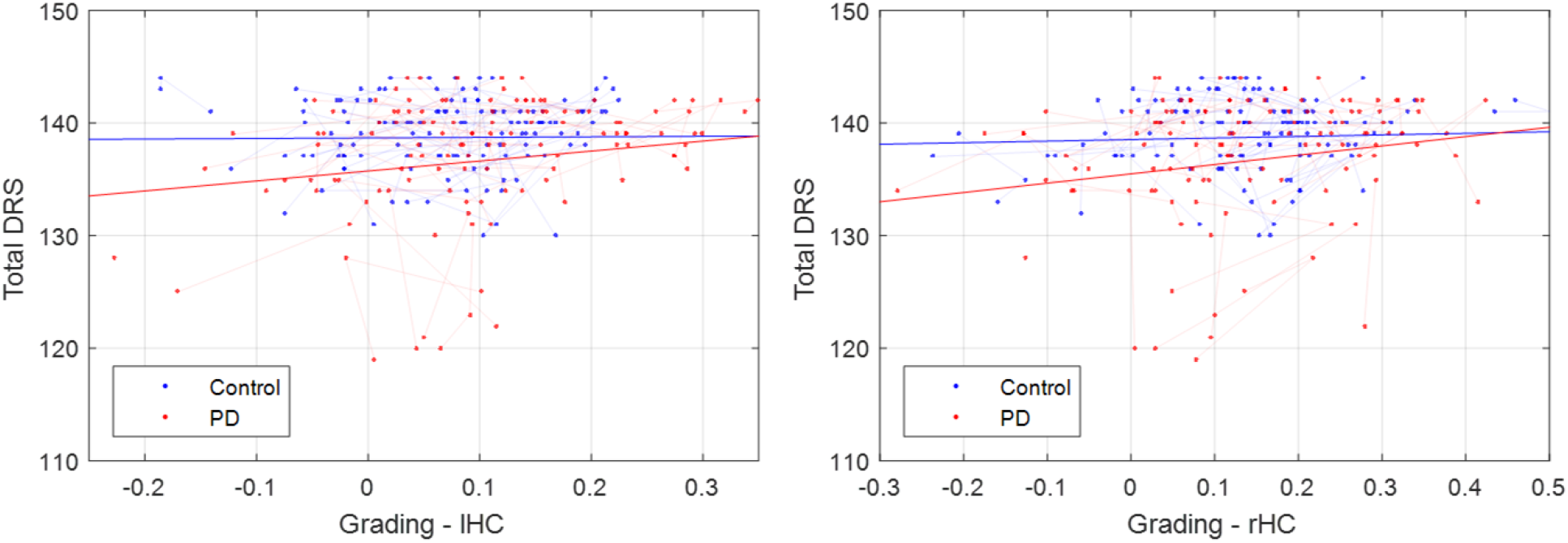
Association between Total DRS and mean SNIPE grading scores in left and right hippocampi. DRS= Dementia Rating Scale. SNIPE= Scoring by Nonlocal Image Patch Estimator. PD= Parkinson’s Disease.

### Total Dementia Rating Scale

Controlling for age and sex, we did not find a significant association between WMH load and Total DRS score in either group. Total DRS significantly decreased with age (t=-2.23, p<0.04) and decrease in hippocampal SNIPE grading in the PD group (t_LeftHC_=2.83, p_LeftHC_=0.005, t_RightHC_=2.55, p_RighHC_=0.01), with a marginal interaction (Figure 4, t_LeftHC_=-1.80, p_RighHC_=0.07, t_LeftHC_=-1.58, p_RighHC_=0.10). We also performed a second level analysis, assessing the associations with individual DRS subscores (Figure S.1, Table S.1). Memory, attention, and initiation preservation subscores showed similar relationships with hippocampal SNIPE grading scores, while conceptualization and constructions were not significantly associated. To ensure that these associations were not just downstream effects of PD-related pathologies, we repeated the analysis, including UPDRSIII, step count in the gait trial, and substantia nigra atrophy as covariates in the models. The obtained results were similar in terms of estimated t statistics and p values. There was no significant impact of sex on the Total DRS measurement in either group.

### Motor Symptoms

Controlling for age and sex, higher WMH load was significantly associated with higher UPDRSIII scores in PD (Figure 5.a, t=3.41, p=0.0008), but not in controls (t=0.26, p=0.79), with a significant interaction (t=2.30, p=0.02). UPDRSIII also significantly increased with age (t=4.75, p<0.00001) and decrease in SN DBM (Figure 5.b-c, t_LeftSN_=-2.02, t_RightSN_=-1.88, p<0.05), with a significantly greater impact in PD (t_LeftSN_=-2.10, t_RightSN_=-1.73, p<0.04). There was no significant impact of sex on the UPDRSIII measurement in either group.

**Figure 5.**
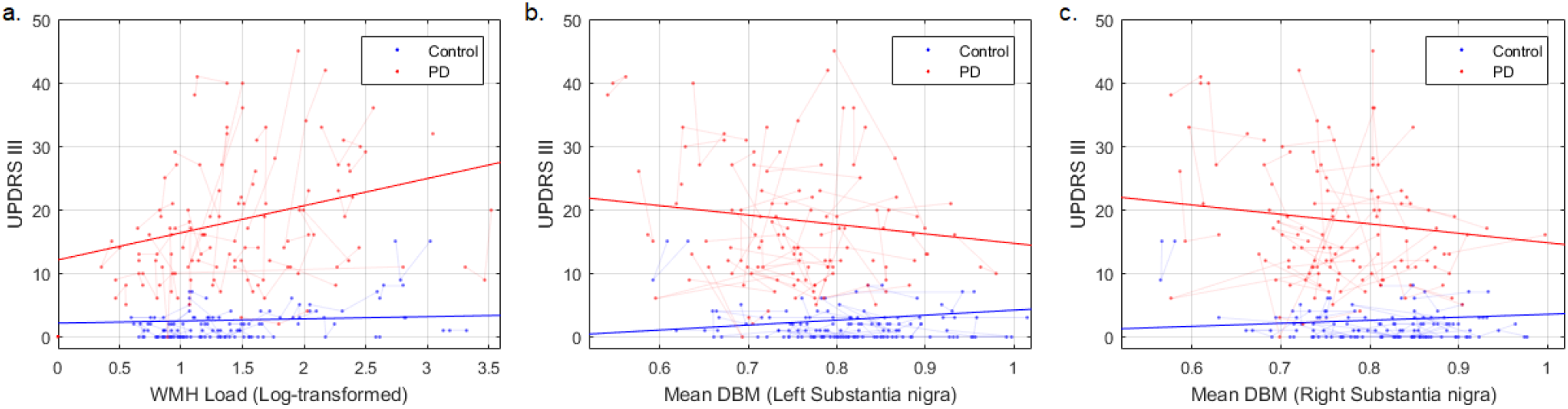
Association between UPDRSIII, WMH burden and mean DBM scores in left and right substantia nigra. UPDRS= Unified Parkinson’s Disease Rating Scale. WMH= White Matter Hyperintensities. DBM= Deformation Based Morphometry.

### Gait

The number of steps in the walking trial significantly increased with age (t=2.50, p=0.01) and increased WMH load (t=2.13, p=0.03) in both groups (Figure 6.a). There was no significant interaction (t=0.95, p=0.34). An increased step count was significantly associated with a decrease in the mean DBM in the right hippocampus in the PD group (Figure 6.c, t=-2.02, p=0.04) with a significant interaction (t=2.18, p=0.03), but not the left (Figure 6.b, p>0.11). Controlling for age and sex, there was no significant association between mean DBM in the substantia nigra and gait in either controls or PD patients (p>0.5). There was no significant impact of sex on gait in either group.

**Figure 6.**
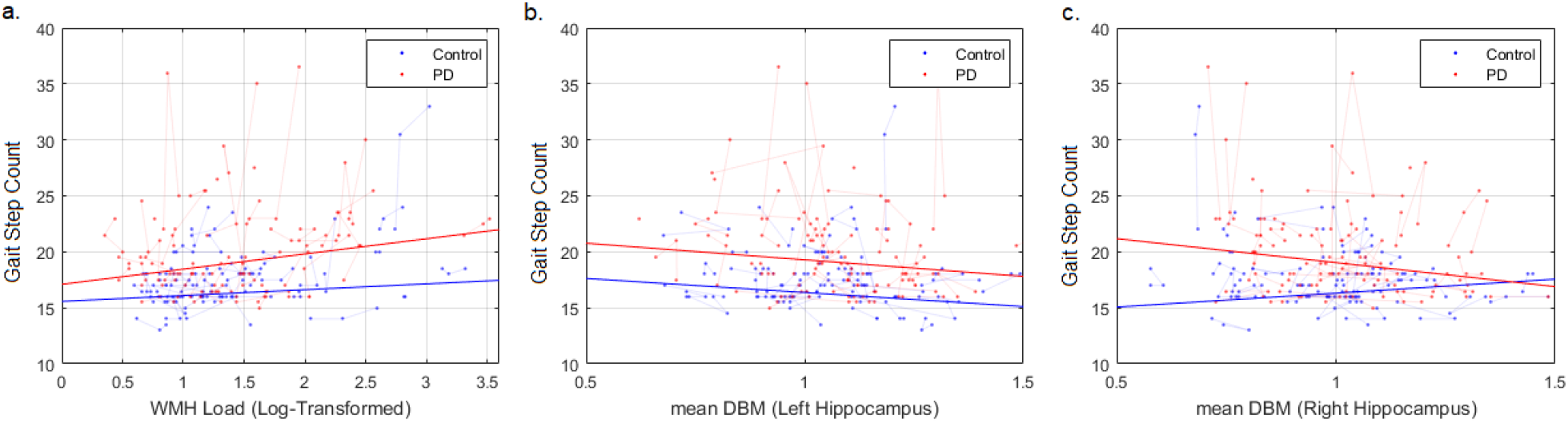
Association between gait, WMH burden and mean DBM scores in left and right hippocampi. WMH= White Matter Hyperintensities. DBM= Deformation Based Morphometry.

## DISCUSSION

In this study, we investigated the impact of grey matter and white matter pathology, in a longitudinal sample of PD patients and age-matched controls. In addition to the overall pattern of increased atrophy associated with age in both controls and PD patients, we found significantly greater bilateral atrophy in the substantia nigra and accumbens areas as well as the right subthalamic nucleus in the PD patients. These findings are in line with the pathologic hallmarks of PD and the previous imaging studies in the literature^3,37^.

Also in line with previous studies, atrophy in the right hippocampus was associated with slowing in gait as well as future cognitive decline in PD patients^5^ (Figures 3.c and 6.c). In addition, bilateral hippocampal atrophy was associated with poorer cognitive performance in both groups, with a marginally greater impact in the PD patients (Figure 4). The hippocampus is involved in spatial memory as well as sensorimotor integration, contributing to both gait and cognitive function. Loss of hippocampal integrity is a well-established contributor to cognitive decline in the aging population. This is also in line with previous studies reporting contribution of Lewy body and possibly coexisting Alzheimer’s disease pathology to dementia in PD patients^2^. Furthermore, to ensure that these associations were not just downstream effects of PD-related pathologies, we repeated the analysis, including UPDRSIII, gait step count, and substantia nigra atrophy as covariates in the models, and obtained similar relationships between total DRS score and hippocampal atrophy.

Consistent with previous findings, PD patients had significantly greater atrophy in the bilateral substantia nigra^3,37^. Substantia nigra atrophy increased with age and was associated with poorer motor perfomance in the PD patients (Figure 3), as also reported in previous studies^3,4^. Substantia nigra contains dopamine neurons whose degeneration accounts for all the key clinical features of PD. The sensitivity of DBM (which is based only on T1w MRI) in detecting substantia nigra atrophy in PD is yet another clinically important finding in this study.

WMH volume increased with age in both controls and PD patients (Figure 2.e). Similar to the majority of previous studies which report no disease-specific differences in WMH loads in PD patients and age-matched normal controls, we did not find a significant difference in WMH burden between controls and PD patients^16,18,38,39^. However, increase in WMH burden was associated with greater cognitive decline and motor and gait deficits, as previously reported in the literature^7,8,16–21^ (Figures 3.a, 5.a, and 6.a, respectively). Taken together, our results support the conclusion that even though PD patients do not present with greater WMH loads than normal aging individuals, comorbid existence of WMHs exacerbates the cognitive and motor sysmptoms in PD by aggravating the already defective neuronal connections.

Few studies have investigated the impact of longitudinal progression of WMHs on cognition in PD. We found that PD patients that were diagnosed with dementia at the last visit (after three years) had greater WMH load increase compared to those that remained stable. This is in line with another previous study, which found periventriventricular WMH progression at 30-month follow-up to be associated with conversion to dementia^40^.

Compared to controls, our PD patients had significantly lower total and LDL cholesterol levels. This might be due to selection bias or better risk management of the PD group receiving tertiary care. The PD group also had significantly higher homocysteine levels. This is in line with previous studies, reporting higher homocysteine levels in PD patients, particularly those treated with L-Dopa^41^.

Higher total cholesterol and LDL cholesterol levels were associated with lower WMH volumes in both controls and PD patients (Table 2). Although surprising, this is in agreement with previous studies showing a positive impact of hyperlipidimia on lowering WMH burden in three independent large samples, suggesting a relatively protective role of hyperlipidimia on small-vessel disease^42,43^. Unlike previous studies, we did not find an association between hypertension and WMH burden^16,42^ (Table 2). This might be due to the relatively smaller sample size or that the blood presure levels were well-controlled in both controls and PD patients in our sample (Table 1). Also, WMH burden was associated with diabetes and higher fasting glucose levels in the control group, but not the PD patients. This might also be due to the small sample size or the patient selection process.

There are several strengths to the current study. Longitudinal MRI and clinical data were consistently acquired from the patients and age-matched controls across all visits. DBM, SNIPE, and WMH measurements were estimated using extensively validated and widely used automated methods ^3,4,11,16,22,35–37^, allowing refined and accurate assessment of the grey matter and white matter pathology in the same population. Relevant medication (both PD and non-PD) was included as covariates in the models. The main limitation of our study concerns its relatively small sample size. It should also be noted that patients with dementia or a history of stroke at baseline were excluded from this study, leading to a select patient group. Replication in larger cohorts are therefore necessary to further confirm these findings.

In conclusion, this study suggests that grey matter atrophy and WM lesions contribute to the cognitive decline and motor deficits in PD. In accordance with previous studies, we found that PD patients had greater bilateral substantia nigra atrophy compared to controls, contributing to poorer motor performance. Hippocampal atrophy also affected both cognition and gait in the PD patients. Together with the evidence from previous studies, our results suggest that even though WMH burden does not differ between PD patients and age-matched otherwise healthy individuals, it aggravates the cognitive and motor symptoms in PD patients. Since many risk factors that are associated with WMHs and their progression are potentially modifiable through treatment and lifestyle changes, the impact of these lesions on cognitive and motor function in PD pinpoints a need for assessment and treatment of these risk factors with a higher priority in PD patients. Given that currently, early intervention and preventive strategies are the key elements in preventing cognitive decline in the aging populations, PD patients might also benefit from systematic cardiovascular risk modification.

## Contributors

MD: Study design, analysis and interpretation of the data, drafting the manuscript.

MG: Data collection and preparation, revision of the manuscript.

AS: Study design, revision of the manuscript.

SD: Study design, interpretation of the data, revision of the manuscript.

RC: Study design, interpretation of the data, revision of the manuscript.

